# Bayesian Optimisation for Neuroimaging Pre-processing in Brain Age Prediction

**DOI:** 10.1101/201061

**Authors:** Jenessa Lancaster, Romy Lorenz, Rob Leech, James H Cole

**Affiliations:** Computational, Cognitive & Clinical Neuroimaging Laboratory, Division of Brain Sciences, Department of Medicine, Imperial College London, London, UK

**Keywords:** brain ageing, Bayesian optimisation, T1-MRI, machine learning, pre-processing

## Abstract

Neuroimaging-based age predictions using machine learning have been shown to relate to cognitive performance, health outcomes and progression of neurodegenerative disease. However, even leading age-prediction algorithms contain measurement error, motivating efforts to improve experimental pipelines. T1-weighted MRI is commonly used for age prediction, and the pre-processing of these scans involves normalisation to a common template and resampling to a common voxel size, followed by spatial smoothing. Resampling parameters are often selected arbitrarily. Here, we sought to improve brain-age prediction accuracy by optimising resampling parameters using Bayesian optimisation.

Using data on N=2001 healthy individuals (aged 16-90 years) we trained support vector machines to i) distinguish between young (<50 years) and old (>50 years) brains and ii) predict chronological age, with accuracy assessed using cross-validation. We also evaluated model generalisability to the Cam-CAN dataset (N=648, aged 18-88 years). Bayesian optimisation was used to identify optimal voxel size and smoothing kernel size for each task. This procedure adaptively samples the parameter space to evaluate accuracy across a range of possible parameters, using independent sub-samples to iteratively assess different parameter combinations to arrive at optimal values.

When distinguishing between young and old brains a classification accuracy of 96.25% was achieved, with voxel size = 11.5mm^3^ and smoothing kernel = 2.3mm. For predicting chronological age, a mean absolute error (MAE) of 5.08 years was achieved, with voxel size = 3.73mm^3^ and smoothing kernel = 3.68mm. This was compared to performance using default values of 1.5mm^3^ and 4mm respectively, which gave a MAE = 5.48 years, a 7.3% improvement. When assessing generalisability, best performance was achieved when applying the entire Bayesian optimisation framework to the new dataset, out-performing the parameters optimised for the initial training dataset.

Our study demonstrates the proof-of-principle that neuroimaging models for brain age prediction can be improved by using Bayesian optimisation to select more appropriate pre-processing parameters. Our results suggest that different parameters are selected and performance improves when optimisation is conducted in specific contexts. This motivates use of optimisation techniques at many different points during the experimental process, which may result in improved statistical sensitivity and reduce opportunities for experimenter-led bias.

## Introduction

The ageing process affects the structure and function on the human brain in a characteristic manner that can be measured using neuroimaging. This quantifiable relationship was key to the early demonstration of the proof-of-principle of voxel-based morphometry (Good et al., 2001) and to this day represents one of the most robust known relationships between a measurable phenomenon (i.e., ageing) and brain structure, making it ideal for evaluating novel neuroimaging analysis tools. More recently, researchers have used this relationship to develop neuroimaging-based tools for predicting chronological age in healthy people using machine learning (Franke et al., 2010; Cole et al., 2017b). A ‘brain-predicted age’ determined from magnetic resonance imaging (MRI) scans represents an intuitive summary measure of the natural deterioration associated with the effects of the ageing process on the brain, and may have the potential to serve as biomarker of age-related brain, or even general, health (Cole, 2017).

The extent to which brain-predicted age is greater than an individual’s chronological age has been associated with accentuated age-associated physical and cognitive decline (Cole et al., 2017c). Specifically, an ‘older’-appearing brain has been associated with decreased fluid intelligence, reduced lung function, weaker grip strength, slower walking speed and an increased likelihood of mortality in older adults (Cole et al., 2017c). Factors which could contribute to an increased brain-predicted age include genetic effects, having sustained a traumatic brain injury, certain neurological or psychiatric conditions, or poor physical health (Koutsouleris et al., 2013; Löwe et al., 2016; Steffener et al., 2016; Cole et al., 2017a; Cole et al., 2017d; Pardoe et al., 2017). Potentially, individuals at increased risk of experiencing the negative consequences of brain ageing, such as cognitive decline and neurodegenerative disease, could be identified by measuring brain-predicted age in clinical groups or even screening the general population.

Despite promising results to-date, models for generating brain-predicted age still continue to contain measurement error, and efforts to improve accuracy and particularly, generalisability, to data from different MRI scanners are warranted. Training on large cohorts of healthy adults gives the lowest mean absolute error (MAE) rates are between 4-5 years (Wang and Pham, 2011; Steffener et al., 2016). Notably, individual errors range across the population, from perfect prediction, to discrepancies as great as 25 years. While brain-predicted age has high test-retest reliability (Cole et al., 2017b), and a proportion of this variation likely reflects underlying population variability, certainly a substantial amount of noise remains. Reducing noise and improving prediction accuracy and generalisability is essential for if such approaches are to be applied to individuals in a clinical setting, the ultimate goal of any putative health-related biomarker.

A key issue in brain-age prediction, along with many other neuroimaging approaches, is the choice of methods for extracting features or summary measures from raw data for further analysis (Jones et al., 2005; Klein et al., 2009; Franke et al., 2010; Andronache et al., 2013; Shen and Sterr, 2013). For example, the majority of brain-age prediction pipelines have used T1-weighted MRI data and either generated voxelwise maps of brain volume (Cole et al., 2015) or summary measures of cortical thickness and subcortical volumes (Liem et al., 2017). The parameters set during the image pre-processing are commonly the defaults supplied by the software developer, or based on prior studies in contrasting study samples. Nevertheless, the choice of pre-processing parameters may have a strong influence on the outcome of any subsequent data modelling, and ideally should be optimised on a case-by-case basis. This optimisation is rarely conducted, as trial-and-error approaches are time-consuming and often ill-posed. Importantly, this issue may reduce experimental precision, which increases the likelihood of false positives and reduces reproducibility. In the worst case scenario, this may encourage p-hacking, whereby pre-processing is manually optimised based on minimising the resultant p-values of the subsequent hypothesis testing. Here, we outline a principled Bayesian optimisation strategy for identifying optimal values for pre-processing parameters in neuroimaging analysis, implementing sub-sampling to avoid bias. We then demonstrate proof-of-principle applied to the problem of age prediction using machine learning.

Bayesian optimisation is an efficient and unbiased approach to the parameter selection problem, which avoids both the failure to adequately search the value space, and the drawbacks of an exhaustive search. Instead, it utilizes a guided sampling strategy to observe a subgroup of points from within the possible parameter space, testing values on subsets of the total subject population (Brochu et al., 2010; Snoek et al., 2012). The data division strategy ensures performance tests always reflect out-of-sample prediction, and always evaluate differing conditions on separate data, reducing the risk of overfitting. Intelligent selection of a small number of points for evaluation allows the characterisation of parameter space and the solution of the optimisation problem to be accomplished in fewer steps (Pelikan et al., 2002).

The current work aimed to use a Bayesian optimisation framework to optimise image pre-processing parameters for: i) distinguishing the brains of young and old adults (classification), ii) predicting chronological age (regression), and iii) evaluating the generalisability of the resulting optima to an independent dataset. We hypothesised that by using Bayesian optimisation we would improve model accuracy compared to previously used ‘non-optimised’ values. The study was designed to show proof-of-principle of the applicability of Bayesian optimisation to help improve neuroimaging pre-processing in a principled and unbiased fashion.

## Materials and Methods

### Neuroimaging Datasets

This study used the Brain-Age Healthy Control (BAHC) dataset, compiled from 14 public sources (see Table 1) and used in our previous research (e.g., Cole et al., 2015). Data included T1-weighted MRI from 2003 healthy individuals aged 16 to 90 years (male/female = 1016/987, mean age = 36.50 ± 18.52). All BAHC participants were either originally included in studies of the healthy population, or as healthy controls from case-control studies and were screened according to local study protocols to ensure they had no history of neurological, psychiatric or major medical conditions. Images were acquired at 1.5T or 3T with standard T1-weighted MRI sequences. Ethical approval and informed consent were obtained locally for each study covering both participation and subsequent data sharing.

**Table 1.** Data sources for healthy brain age training sample

| Cohort | N | Age<br>mean (SD) | Age<br>range | Sex<br>male/female | Repository details | Scanner<br>(Field strength) | Scan | Voxel dimensions |
| --- | --- | --- | --- | --- | --- | --- | --- | --- |
| ABIDE (Autism Brain Imaging Data Exchange) | 184 | 25.93 (6.66) | 18-48 | 161/23 | INDI | Various (all 3T) | MPRAGE | Various |
| Beijing Normal University | 181 | 21.25 (1.92) | 16-28 | 72/109 | INDI | Siemens (3T) | MPRAGE | 1.33x1.0x1.0 |
| Berlin School of Brain & Mind | 49 | 30.99 (7.08) | 20-60 | 24/25 | INDI | Siemens Tim Trio (3T) | MPRAGE | 1.0x1.0x1.0 |
| CADDementia | 12 | 62.33 (6.26) | 55-79 | 9/3 | http://caddementia.grand-challenge.org | GE Signa (3T) | 3D IR-FSPGR | 0.9x0.9x1.0 |
| Cleveland Clinic | 31 | 43.55<br>(11.14) | 24-60 | 11/20 | INDI | Siemens Tim Trio (3T) | MPRAGE | 2.0x1.0x1.2 |
| ICBM (International Consortium for Brain Mapping) | 322 | 24.84 (5.14) | 24-60 | 177/142 | LONI IDA | Siemens Magnetom (1.5T) | MPRAGE | 1.0x1.0x1.0 |
| IXI (Information eXtraction from Images) | 561 | 48.62<br>(16.49) | 20-86 | 250/311 | http://biomedic.doc.ic.ac.uk/brain-development | Philips Intera (3T);<br>Philips Gyroscan Intera (1.5T);<br>GE Signa (1.5T) | T1-FFE;<br>MPRAGE | 0.9375x0.9375x1.2 |
| MCIC (MIND Clinical Imaging Consortium) | 93 | 32.49<br>(11.95) | 18-60 | 64/29 | COINS | Siemens Sonata/Trio (1.5/3T);<br>GE Signa (1.5T) | MPRAGE;<br>SPGR | 0.625x0.625x1.5 |
| MIRIAD (Minimal Interval Resonance Imaging in Alzheimer's Disease) | 23 | 69.66 (7.18) | 58-85 | 12/11 | https://www.ucl.ac.uk/drc/research/miriad-scan-database | GE Signa (1.5T) | 3D IR-FSPGR | 0.9375x0.9375x1.5 |
| NEO2012 (Adelstein, 2011) | 39 | 29.59 (8.38) | 20-49 | 18/21 | INDI | Siemens Allegra (3T) | MPRAGE | 1.0x1.0x1.0 |
| Nathan Kline Institute (NKI) / Rockland | 160 | 41.49<br>(18.08) | 18-85 | 96/64 | INDI | Siemens Tim Trio (3T) | MPRAGE | 1.0x1.0x1.0 |
| OASIS (Open Access Series of Imaging Studies) | 288 | 44.06<br>(23.04) | 18-90 | 106/188 | http://www.oasis-brains.org/ | Siemens Vision (1.5T)* | MPRAGE | 1.0x1.0x1.25 |
| WUSL (Power, 2012) | 24 | 23.04 (1.42) | 20-24 | 4/20 | INDI | Siemens Tim Trio (3T) | MPRAGE | 1.0x1.0x1.0 |
| TRAIN-39 | 36 | 22.67 (2.56) | 18-28 | 11/25 | INDI | Siemens Allegra (3T) | MPRAGE | 1.33x1.33x1.3 |
| Training set total | 2003 | 36.50<br>(18.52) | 16-90 | 1016/987 | - | - | - | - |

INDI = International Neuroimaging Data-sharing Initiative (http://fcon_1000.projects.nitrc.org) COINS = Collaborative Informatics and Neuroimaging Suite (http://coins.mrn.org) LONI = Laboratory of Neuro Imaging Image & Data Archive (https://ida.loni.usc.edu) ABIDE consortiums comprising data from various sites with different scanners/parameters \*OASIS scans were acquired four times and then averaged to increase signal-to-noise ratio.

Additionally, the Cambridge Centre for Aging Neuroscience (Cam-CAN) neuroimaging cohort was used as an independent validation dataset (Shafto et al., 2014). The Cam-CAN cohort consisted of T1-weighted images from 648 participants aged 18 to 88 years (male/female = 324/324, mean age 54.28 ± 18.56). This study used similar exclusion criteria, including only healthy individuals. MRI scans were T1-weighted and were acquired at 3T. Again ethical approval for Cam-CAN was obtained locally, including the permission to subsequently make anonymised versions of the data publicly-available.

### Pre-processing to Prediction

Normalized brain volume maps were created following the protocol described in (Cole et al., 2015). This involved segmentation of raw T1-weighted images into grey matter maps using SPM12 (University College London, London, UK). Images were normalized to a study-specific template in MNI152 space using DARTEL for non-linear registration (Ashburner, 2007). This step involved resampling to a common voxel size, modulation to retain volumetric information and spatial smoothing; the specific voxel size and smoothing kernel size parameters were chosen by the Bayesian optimisation protocol as detailed below.

After pre-processing, images were converted to vectors of ASCII-format intensity values. These were used as the input features for subsequent classification or regression analysis. This was performed in MATLAB using the support vector machine (SVM) program. For the binary classification problem of predicting younger from older participants, SVMs were used. For predicting age as a continuous variable, SVM regression (SVR) was used, using participants from the full age range (16-90 years). Both SVM and SVR procedures used a linear kernel to map the input data into a computationally-efficient feature space.

### Bayesian Optimisation

Bayesian optimisation was used to identify optimal pre-processing parameters, based on the accuracy of the subsequent model predictions (either classification or regression). Hence, the Bayesian optimisation procedure can be seen as an additional outer layer of analysis, that surrounds the standard pipeline (pre-processing through to model accuracy evaluation). The Bayesian optimisation process runs multiple iterations of this internal pipeline, exploring the parameter space to select varying image pre-processing options based on their influence on the objective function (i.e., classification or regression accuracy).

A key advantage of Bayesian optimisation derives from its ‘surrogate’ model that represents relationship between an algorithm's and the currently unknown objective function. This surrogate model is progressively refined in a closed-loop manner, by automatically selecting points in the parameter space, in order to provide informed coverage of that based, based on the performance of previously sampled points. This aspect makes Bayesian optimisation highly efficient, reducing the number of iterations necessary to identify optima of complex objective functions (Brochu et al., 2010; Lorenz et al., 2017). We used the MATLAB Bayesian Optimization Algorithm (https://uk.mathworks.com/help/stats/bayesian-optimization-algorithm.html) implementation, which internally defines a number of optimisation parameters, including selecting the covariance kernel and tuning the hyper-parameters of the process.

In this study, we aimed to optimise the final normalised voxel size and the full-width half-maximum (FWHM) of the spatial smoothing kernel used during final resampling. Conventionally, these are set at 1.5mm^3^ and either 4mm or 8mm FWHM, respectively for VBM studies conducted using SPM. Using Bayesian optimisation, the range of considered values was extended from common practices. A range of [1,30] mm isotropic was permitted for voxel size and [1,20] mm FWHM for smoothing kernel size. In subsequent analyses with a constant voxel size or smoothing value, the value was informed by this result. Following this test, the search range was reduced to [1,15] and [1,10] for voxel size and smoothing kernel size.

The following examples illustrate the use of Bayesian optimisation to visualize the parameter space and its optima across continuous and categorical variables. Due to a large population and straightforward design, the binary classification problem was expected to perform well and behave in a stable manner. These characterization experiments were therefore carried out in context of classification accuracy.

### Classifying Young and Old Adults

We defined the 500 oldest individuals (aged 51 to 90 years) and the 500 youngest (aged 16 to 22 years) as the “old” and “young” groups for classification. Each iteration of Bayesian optimisation used a subsample of the total subject set, *N* = 1000, to test a combination of pre-processing parameter values. Participants were divided into subsets of size *n* stratified by age, such that each subset was approximately representative distribution of participants from across the age range, resulting in a total of *N/n* iterations. We used *n* = 80 total (40 participants from each group) as a sample for each iteration, giving 1000/80 = 12 iterations of Bayesian optimisation. This included a burn-in phase (i.e., preliminary phase of unevaluated samples to initialise the process) of 5 initial, randomly-sampled points from within the parameter ranges to begin characterization of the search space, followed by 7 iterations of ‘guided’ active sampling. In each iteration a voxel-size and smoothing kernel size combination was selected and used for resampling during DARTEL normalisation of each subject’s images. Normalised images were then converted to feature vectors and a binary classifier was trained and assessed using 10-fold cross-validation. Classifier accuracy was the objective function to be minimised. Bayesian optimisation used the Expected Improvement Plus (EI+) acquisition function, with the default exploration-exploitation ratio of 0.5.

### Regression Prediction of Chronological Age

Next, we used Bayesian optimisation to assess regression models of healthy brain ageing that allows accurate prediction of age in new datasets. This was done by first identifying optimal parameters through Bayesian optimisation, then applying them to the full training dataset and comparing the resulting prediction accuracy to that achieved in the current literature. Here, participant age was used as the outcome (i.e., dependent) variable the vector of brain volume values as the predictors (i.e., independent variables).

Regression analysis also used *n* = 80 participants per iteration, with age values spanning the full range, and with the same Bayesian optimisation hyper-parameters. The MAE in age prediction across 10-fold cross-validation using SVM regression was the objective function to be minimized. To enable both the optimisation search and make use of the full sample size available for training a generalisable regression model, the population was divided into a training set and a held-out test set. Bayesian optimisation was first carried out using 1803 of the 2003 total participants to determine the optimal voxel size and smoothing kernel size values (allowing 22 total optimisation iterations). A regression model was then trained on these 1803 images pre-processed using the identified optimal values, and tested on the 200 held-out participants in the test set (pre-processed using the same optimised parameters) to give an unbiased out-of-sample measure of age prediction performance.

A final regression model was trained on all 2003 participants (the full model) for application to independent datasets. This allowed as to evaluate how well the optimized pre-processing parameters generalized. To achieve this, we tested the full model on the Cam-CAN dataset for independent validation. We compared three possible approaches for this independent validation step: 1) application of the BAHC-derived full model to age prediction on Cam-CAN data, 2) application of the entire Bayesian optimisation framework to the Cam-CAN dataset followed by regression training, 3) using the BAHC-derived pre-processing parameters to process the Cam-CAN dataset, but training a new regression model for age prediction. In case #1 the optimised voxel size and smoothing kernel size from the BAHC dataset were applied to the Cam-CAN dataset. In case #2, the pre-processing were optimized afresh, using only the Cam-CAN dataset. Case #3 is something of an intermediate iteration, generalizing the optimized pre-processing parameters, but not the trained regression model.

### Performance Stability

Finally, we performed several experiments to assess reproducibility and variability of different solutions to the classification task (i.e., young vs. old). This was done to allow inference regarding which image pre-processing parameters had the greatest impact on prediction accuracy, and to establish robustness of the model. We tested the consistency of model solutions across repetitions and participant sets, using different random seeds to create shuffled groups and burn-in points. We also varied acquisition function of the Bayesian approach. This included comparing the results using six different acquisition functions: Expected Improvement (EI), EI per second, EI+, EI per second +, Lower Confidence Bound, and Probability of Improvement. Finally model solutions were compared across differing values for the exploration-exploitation ratio, ranging from 10% to 90% exploration. These tests were conducted in context of the classification accuracy problem, and used voxel-smoothing kernel size ranges of [1,15] and [1,10], respectively; the Expected Improvement Plus (EI+) acquisition function; and an exploration ratio of 0.5.

## Results

### Classification Analyses

Optimised model performance was an accuracy of 96.25% for correct classification of neuroimaging data as either young or old, at a voxel size of 11.5mm^3^ and smoothing kernel size of 2.3mm. The parameter space exploring the expanded range of voxel size and smoothing kernel size values yielded the model shown in Figure 1.

**Figure 1.**
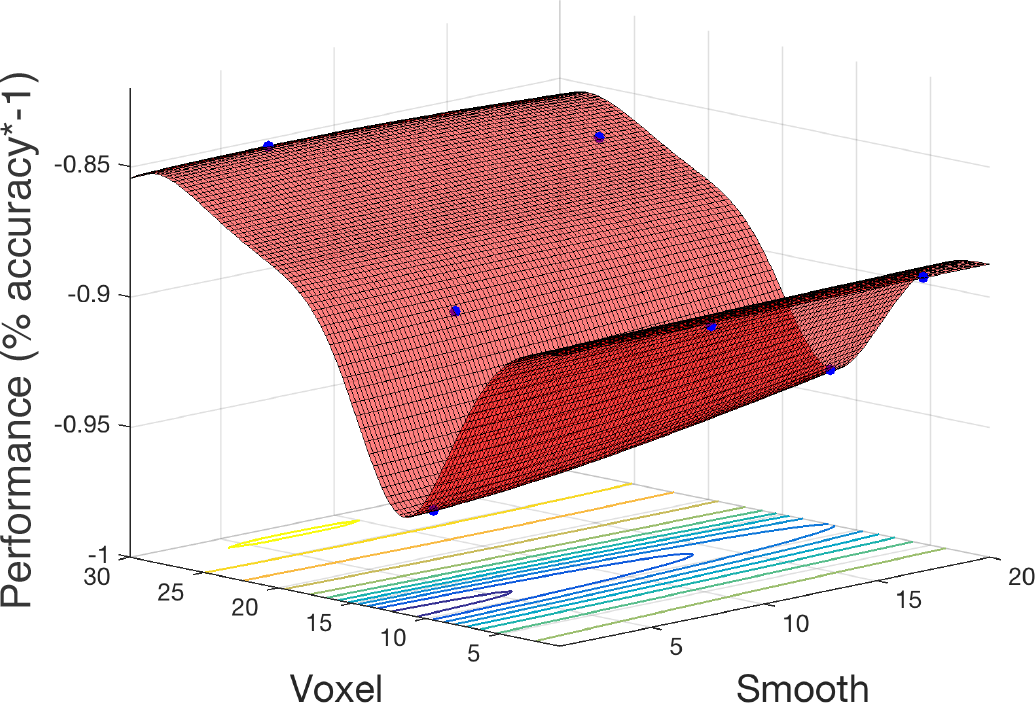
Objective function model for classification of young and old. Objective function model showing the space of voxel size and smoothing kernel values in terms of old versus young classification performance with a linear kernel SVM. The red surface is the function model and represents performance at each location. Points where the parameter space was sampled are shown (blue).

### Age Prediction Regression

Figure 2 shows the objective function model created during regression, when varying smoothing kernel and voxel size ([1:10], [1:15]), and minimizing the MAE in prediction of exact age by Support Vector Regression. The minimum error observed was 7.17 years, and the objective model estimated a minimum error of 8.52 years. The estimated optimal voxel size and smoothing kernel size values were (3.73, 3.68). Following optimization these values were used to pre-process the full dataset, and train a regression model.

**Figure 2.**
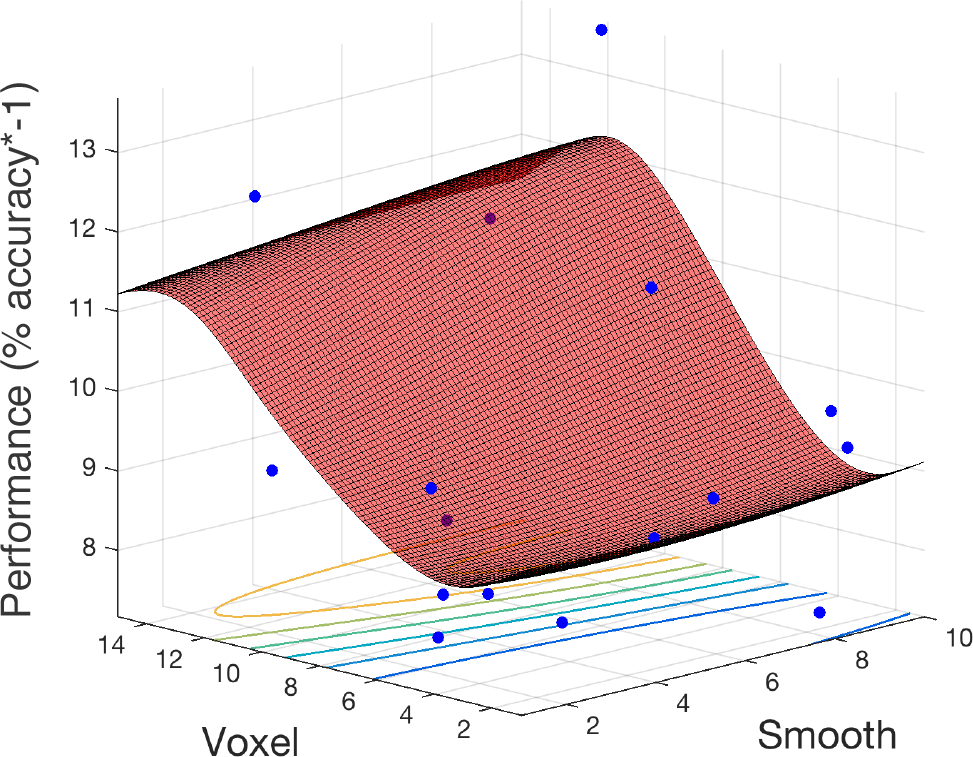
Objective function model for age prediction. Objective function model resulting from optimizing during regression for exact age prediction. The red surface is the function model and represents performance at each location. Points where the parameter space was sampled are shown (blue).

The final resulting model was applied to predict ages for the remaining 200 holdouts and achieved a MAE of 5.08 years. This was compared to MAE = 5.48 years when using the un-optimised pre-processing values. The absolute error observed in any single subject ranged from 0-22.78 years. Figure 3 shows the relationship between predicted age and chronological age for each dataset: the 200 holdout test cases from the BAHC dataset (Fig. 3a). For the BAHC holdout cases, the Pearson’s correlation between predicted and true age was *r* = 0.941, with R^2^ = 0.89 using optimised pre-processing. Using un-optimised pre-processing parameters: *r* = 0.927, R^2^ = 0.86.

**Figure 3.**
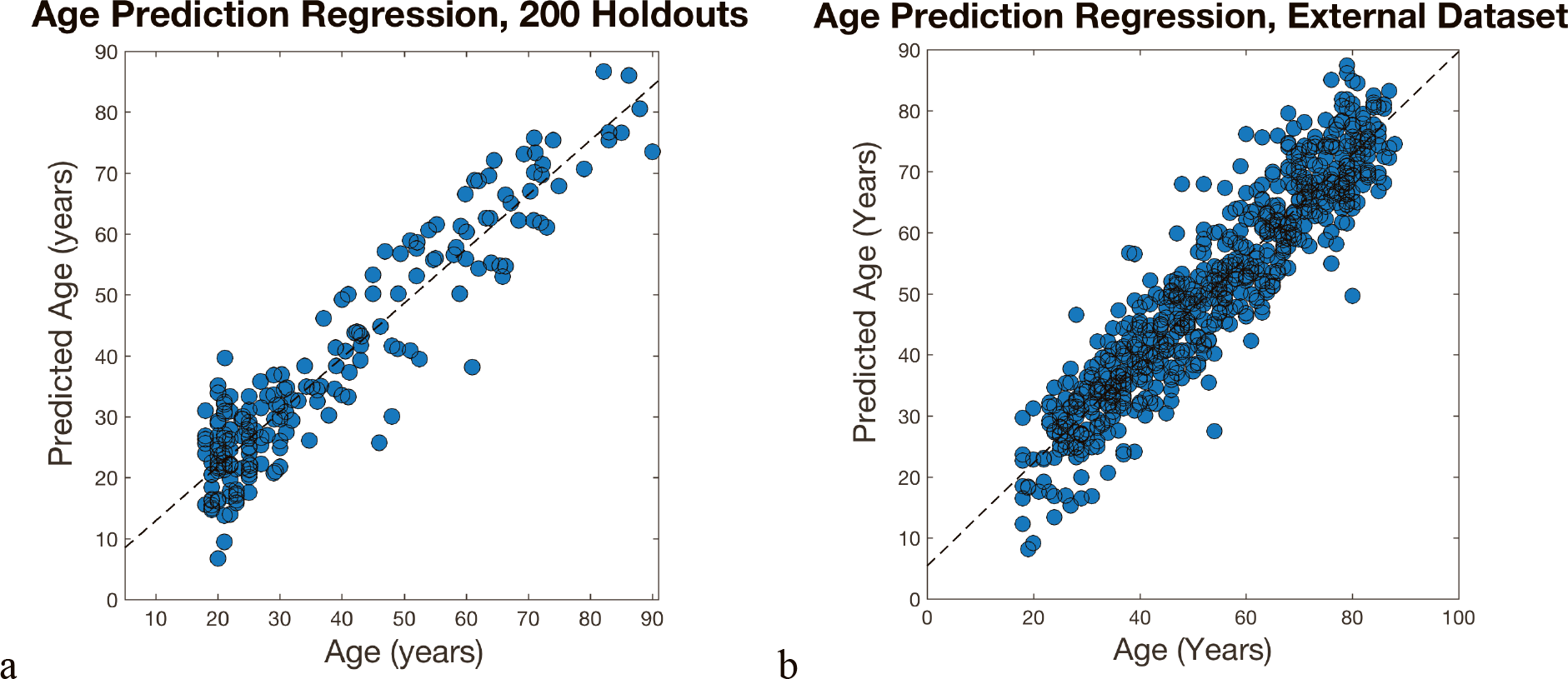
Relationship between chronological age and brain-predicted age. Chronological age (x-axis) plotted against brain-predicted age (y-axis) when testing the BAHC-trained model on (a) the hold-out N=200 test set from BAHC, and (b) on the full Cam-CAN data.

The model generated in the first study of regression was applied to the Cam-CAN dataset in three different ways. 1) The BAHC-trained model was applied to the Cam-CAN data pre-processed with the BAHC-informed optimum voxel size and smoothing kernel size values. This achieved a MAE of 6.08 years, *r* = 0.929, with R^2^ = 0.86 (Figure 3b). This was an improvement compared to the performance when using un-optimised values which had a MAE = 6.76 years. 2) The Cam-CAN data was analysed entirely independently; the full Bayesian optimization framework was instead applied to the Cam-CAN data to discover new, Cam-CAN-specific pre-processing optima, and a new regression model was trained with 588 participants and tested on 60 participants (giving a similar training-testing ratio as used in the BAHC dataset). This yielded a MAE of 5.18 years. 3) the Cam-CAN cohort was pre-processed using the BAHC-informed optimum values but a new regression model was trained and tested within the Cam-CAN set. This model resulted in a prediction error of 6.38 years.

### Classification Model Stability

Stability and reproducibility of model solutions was explored via the classification accuracy problem. Correlation of models in different scenarios are shown in Figure 4. These were; a) across 10 repetitions of the final classification protocol, b) produced by the use of each of six different acquisition functions, and c) using 5 different exploration-exploitation ratios ranging between 90% exploration and 90% exploitation. In all three cases, model solutions showed high cross-correlation across replications, as well as across varying settings of the optimisation process. The choice of acquisition function for Bayesian sampling and the choice of exploration-exploitation ratio of this function had little impact on the final model performance. Similar models were reproducible across repetitions and in randomly-shuffled participant sub-sets. This behaviour implied a stable model exists in the outlined parameter space. These observations support our use of the default acquisition function options for the classification analysis: Expected Improvement Plus (EI+), with an exploration ratio of 0.5.

**Figure 4.**
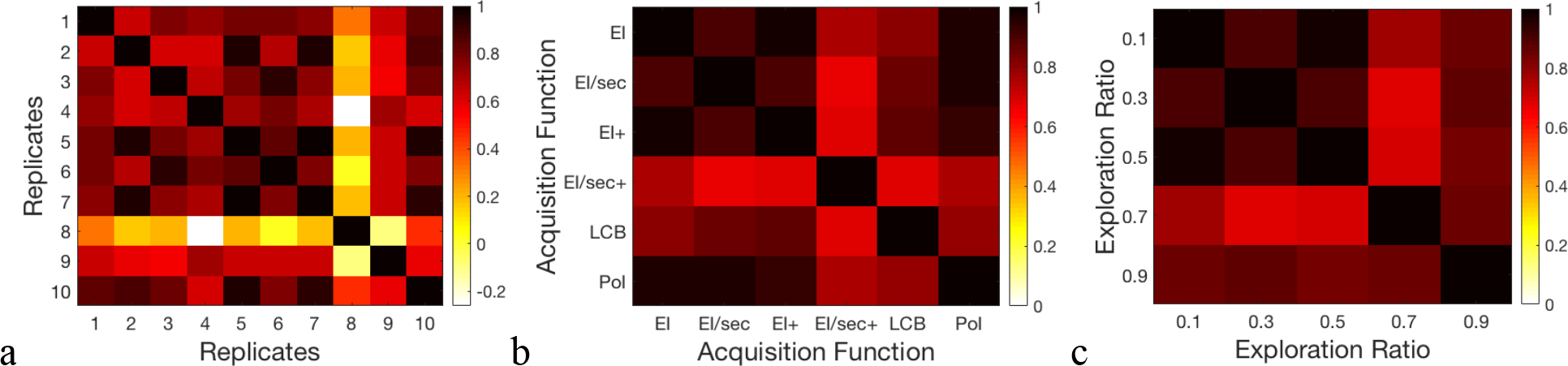
Classification model stability over varying Bayesian optimisation parameters. Correlation heat maps illustrating the relationships between model objective function solutions across (a) replicates of the protocol, (b) six different acquisition functions, and (c) a range of exploration-exploitation ratios between 0.1 and 0.9.

## Discussion

Using Bayesian optimisation, we present a conceptual and practical improvement to conventional pipelines for distinguishing young and old brains or predicting age using neuroimaging data. The Bayesian Optimisation-derived optima for voxel size and smoothing kernel size showed moderate improvement in model performance over ‘un-optimised’ defaults used previously, suggesting that it would be beneficial to incorporate such a process into future ‘brain-predicted age’ research. Our results are important as they suggest that the same pre-processing parameters are not optimal for different prediction tasks (i.e., classification vs. regression) or for different datasets (BAHC vs. Cam-CAN). Often, researchers will apply parameters used in one context to another. This may not necessarily be best practice, and our work shows proof-of-principle that Bayesian Optimisation can be used to improve image pre-processing in a principled and unbiased fashion.

Beyond optimising performance, our Bayesian optimisation approach also allows for relative comparison of the relative influence of different parameters. This potentially provides novel information regarding the prediction problem at hand. For example, here we found that varying voxel size had a much greater impact on overall performance than did smoothing kernel size. This was seen in all experiments; the change in performance across the full range of values was much smaller for smoothing kernel size than voxel size, and is clearly seen in the surface plots (Figs. 1, 2, 4). This suggests that in future neuroimaging pre-processing pipeline design, there is more to be gained from optimising voxel size, rather than smoothing kernel size. The target voxel size for normalisation is often not considered, though has an important impact on the degree of partial volume effects, number of simultaneous statistical tests undertaken, spatial resolution and subsequent inferences made about anatomical specificity. Our findings suggest that more weight needs to be placed on this important parameter when relating volumetric MRI data to age.

Importantly, the conclusions regarding specific optimal values are related to the particular application in which they are tested. Within this study, we observed a notable difference in the optimal voxel size for classification (11.5mm^3^) compared to regression (3.7mm^3^). Potentially, the more gross distinction between young and old brains benefits from a coarser resolution which increases signal-to-noise ratio, while the more subtle patterns underlying gradual age-associated changes in brain structure requires finer-grained representation. Alternatively, the much larger voxel size identified here could result in better classification by reducing data dimensionality, with this size representing the optimal trade-off between representing the information and simplifying a crowded feature space for more effective classification. Either way, the discrepancy in optimal voxel size between classification and regression reinforces the point that systematic evaluation of parameter specifications should be conducted case-by-case. Commonly, ‘one-size-fits-all’ is the prevailing heuristic for setting pre-processing parameters in neuroimaging analysis, where the defaults are often assumed to be adequate. Our findings show that this is not the case, supporting the use of optimisation techniques, to improve experimental precision.

The age-prediction accuracy achieved in the BAHC dataset (MAE = 5.08 years) is comparable to the current performance seen in similar research (Wang and Pham, 2011; Konukoglu et al., 2013; Mwangi et al., 2013; Irimia et al., 2014; Cole et al., 2015; Steffener et al., 2016; Cole et al., 2017b; Cole et al., 2017c). In these studies, pre-processing parameters are set somewhat arbitrarily (c.f. Franke et al., 2010). Here, Bayesian optimisation method offered a more principled approach to this level of accuracy. The prediction tools used here were common methods selected for computational efficiency (i.e., SVMs), in contrast with some of the studies capitalising on state-of-the-art techniques and advanced modelling such as deep learning or Gaussian process regression (Cole et al., 2017b; Cole et al., 2017c). One limitation of Bayesian optimisation is the computational time needed to derive optima, which may slow its adoption with newer, computationally-intensive algorithm. The duration of the optimisation also depends on the process to be optimised. Here we selected image resampling parameters. One important target for optimisation should be image registration, however the speed of most non-linear registration algorithms currently makes this a time-consuming goal.

We explored the generalisability of the model and the general framework. The resulting MAE in the Cam-CAN dataset of 6.08 years (BAHC-defined model and parameters), 5.18 years (Cam-CAN defined model and parameters) and 6.38 years (BAHC-defined parameters, Cam-CAN model) provides some important insights. The BAHC-derived model produced reasonable performance, the highest accuracy was achieved when re-optimising and re-training within the independent cohort, using the full Bayesian optimisation framework. This suggests that there is unlikely to a be ground-truth optimum for a given parameter, highlighting the importance of defining such optima in a given context. This leads to another limitation of the approach, that is sufficiently large datasets are necessary for the optimisation to work effectively. While this is increasing possible due to the drive to share data and the availability of large, publicly-accessible cohorts (e.g., Alzheimer's Disease Neuroimaging Initiative, Human Connectome Project, UK Biobank), this may not be possible in certain clinical contexts, particularly regarding rarer diseases. Nevertheless, the ‘generalised’ performance of the model was still reasonable in this case, which suggests that it is incumbent on researchers to decide what constitutes sufficient prediction accuracy in each context.

Bayesian optimisation is a robust and elegant way to tune pre-processing pipelines in an efficient and automated manner. In addition to the parameter optima, an additional strict, unbiased estimate for performance and generalisability is generated. The objective function model created provides detailed information on model performance and new insights can be gained from mapping the entire parameter space by enabling visualization of relationships between key components of the analysis and performance. This could allow for informed decision-making in experimental design, such as allowing for cost-benefit analysis in the case where optimal parameters only lead to marginal improvement in performance over other values which are easier, quicker, or less costly to enact. This could be critical in applications where the varied inputs represent expensive or invasive procedures, such as MRI scanning or obtaining CSF samples from lumbar punctures.

Though our analysis focused on neuroimaging pre-processing to illustrate the strengths of Bayesian optimisation methods, the potential applications are far-reaching. Optimisation could be applied anywhere in the experimental pipeline: in questions of experimental design, stimuli choice, data acquisition, statistical method or algorithm selection, prediction methods or final model selection. The current literature on Bayesian optimisation topics applies mainly to tuning of machine learning algorithms (Snoek et al., 2012) and though machine learning is widely used in neuroscience, few studies have capitalized on this strategy to improve neuroimaging analysis or neuroimaging-based prediction (Lorenz et al., 2016; Lorenz et al., 2017). In machine learning contexts, and especially in applied multi-disciplinary fields like neuroscience, where researchers may not necessarily have expertise regarding every relevant experimental parameter, more widespread use of *a priori* unbiased parameter optimisation could be greatly beneficial.

Our study shows the value of Bayesian optimisation to improve neuroimaging pre-processing for estimate brain-predicted age, a potential biomarker of healthy brain ageing. Future research into brain ageing and other neuroscientific areas could benefit from applying principled optimisation approaches to improve study sensitivity and reduce bias.

## Author contributions

RoLe and JC conceived and designed the study. RoLo, RoLe, JC developed the methods. JL analysed and interpreted data. RoLe and JC interpreted data. JL, RoLo, RoLe and JC drafted and revised the manuscript.

## Acknowledgements

The authors would like to acknowledge the principal investigators of the studies who made their data openly accessible for research.

